## Supplemental Information for "TRPγ regulates lipid metabolism through *Dh44* neuroendocrine cells"

**This PDF File includes:**

Figure 2-figure supplement 1, Figure 3–figure supplement 1, Figure 5–figure supplement 1, Figure 6-figure supplement 1 and Figure 7–figure supplement 1.

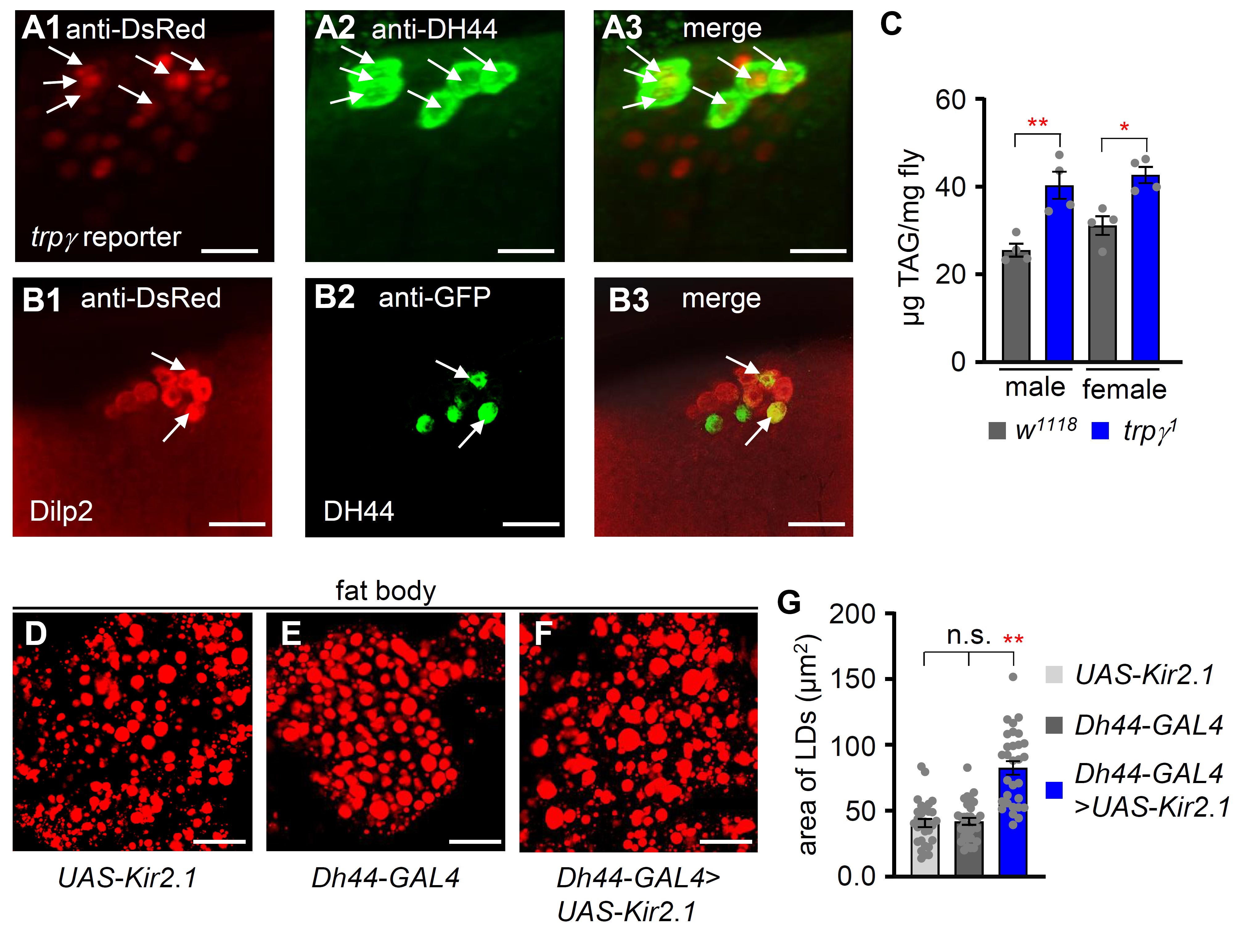
 **Figure 2–figure supplement 1:** Expression of *trpγ*, *Dh44* and *dilp2* in the brain of *Drosophila*. (**A1**–**A3**) Immunohistochemistry of brain tissue in adult flies. **A1**, anti-DsRed staining of *trpγ* reporter (*UAS*-*dsred*;*trpγ^G4^*). **A2**, anti-DH44 staining of *Dh44* cells in the brain. **A3**, merged expression of A1 and A2. (**B1**–**B3**) Immunohistochemistry of brain tissues (*UAS*-*mCD8*::*GFP*/+;*dilp2*-*mcherry*/*Dh44*-*GAL4*). **B1**, Anti-DsRed staining of *dilp2* reporter. **B2**, anti-GFP staining of *Dh44* cells. **B3**, merged expression of B1 and B2. Arrows indicate the co-expression of *dilp2* and DH44 in two neurons. (**C**) Total TAG levels in the whole-body extracts obtained from males and females, separately (n=4). (**D**–**F**) Fat body stains with Nile red. **D** is from the fat body of *UAS-Kir2.1,* **E** is from *Dh44-GAL4* flies, and **F** is from flies with *Dh44* neurons ablated with *UAS*-*Kir2.1.* Scale bar represents 50 µm. (**G**) Measurement of area of LDs in *UAS*-*Kir2.1*, *Dh44*-*GAL4*, and *Dh44*-*GAL4>UAS*-*Kir2.1*, respectively (n=3).

Means ±SEMs. Single factor ANOVA with Scheffe’s analysis was used as a *post hoc* test to compare multiple sets of data. The asterisks indicate significance from control (^*^*P* < 0.05, ^**^*P* < 0.01). Each dot indicates distribution of individual sample values.

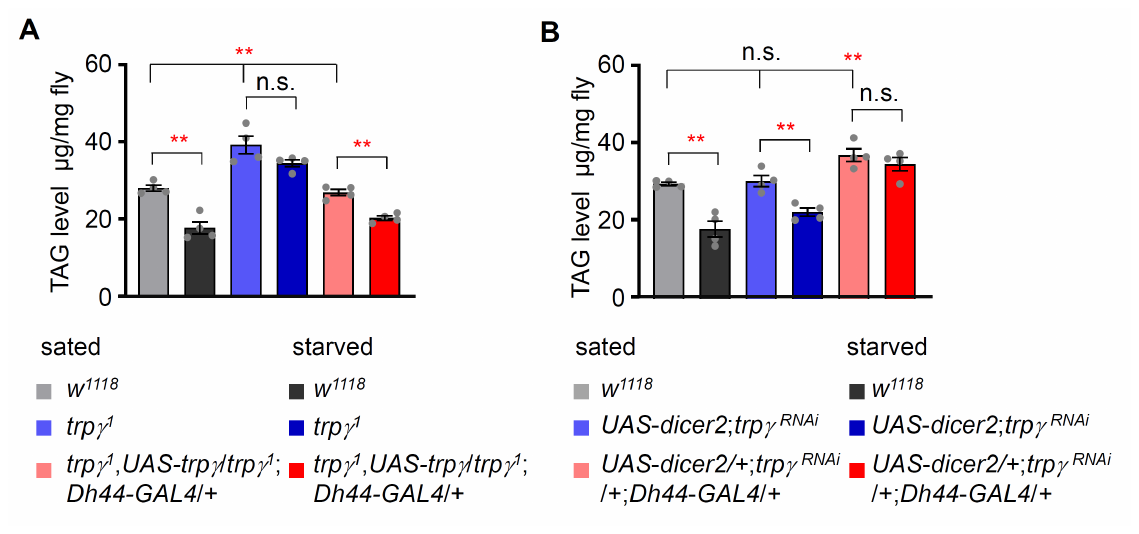
**Figure 3–figure supplement 1:** Measurement of TAG levels after overexpressing *UAS*-*trpγ* and knock-down of *trpγ^RNAi^* in *Dh44* neurons. (**A**) Measurement of total TAG levels in *w^1118^*, *trpγ^1^,* and *Dh44*-*GAL4*/*UAS*-*trpγ* under sated and starved conditions (n=4). (**B**) Measurement of total TAG level in *w^1118^*, *UAS-dicer2;trpγ^RNAi^*, and *UAS-dicer2/+; trpγ^RNA^*^i^/+;*Dh44*-*GAL4*/*+* under sated and starved conditions (n=4).

Means ±SEMs. Single factor ANOVA with Scheffe’s analysis was used as a *post hoc* test to compare multiple sets of data. The asterisks indicate significance from control (^*^*P* < 0.05, ^**^*P* < 0.01). Each dot indicates distribution of individual sample values.

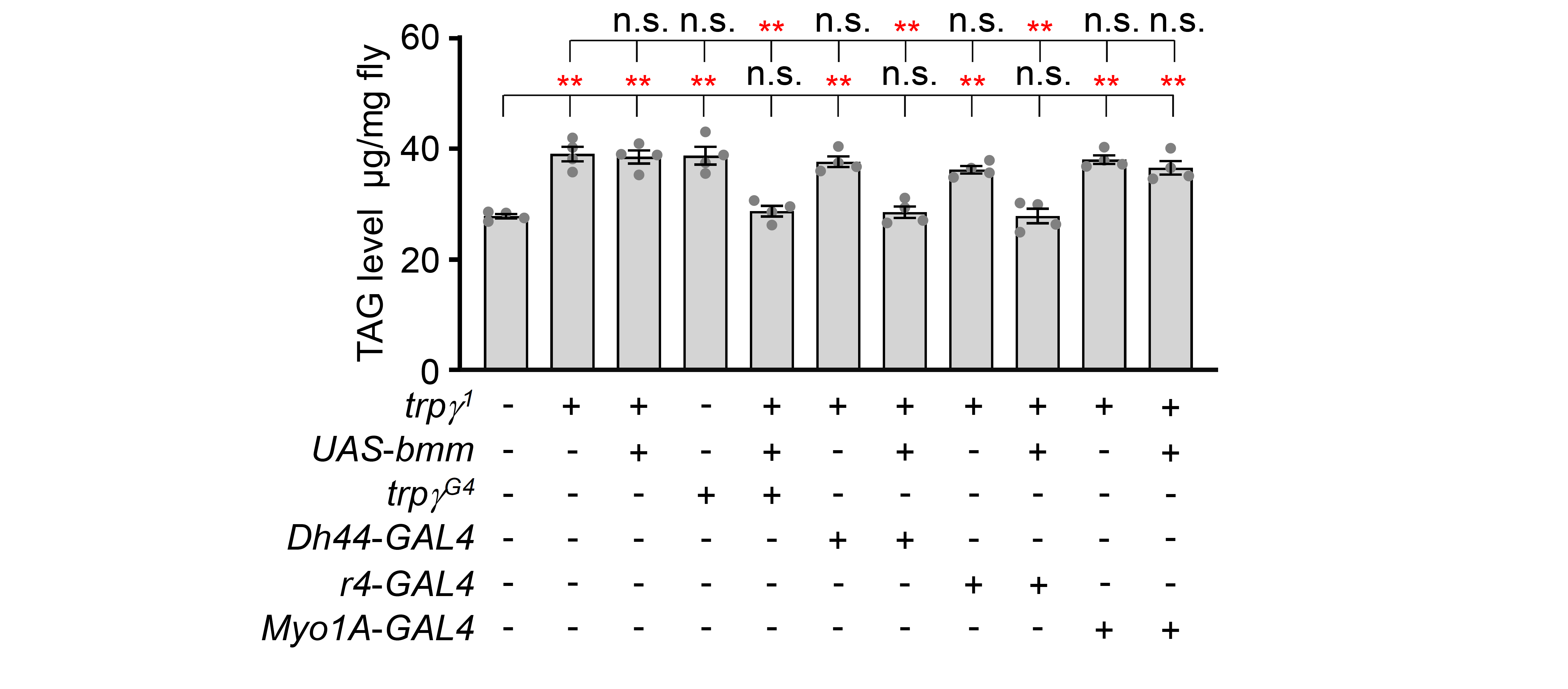

**Figure 5–figure supplement 1:** Measurement of TAG level after expression of *UAS*-*bmm* in *trpγ^G4^*, *Dh44*-*GAL4*, *r4*-*GAL4*, and *Myo1A*-*GAL4* (n=4).

Means ±SEMs. Single factor ANOVA with Student t-test was used as a *post hoc* test to compare data. The asterisks indicate significance from the controls (^**^*P* < 0.01).

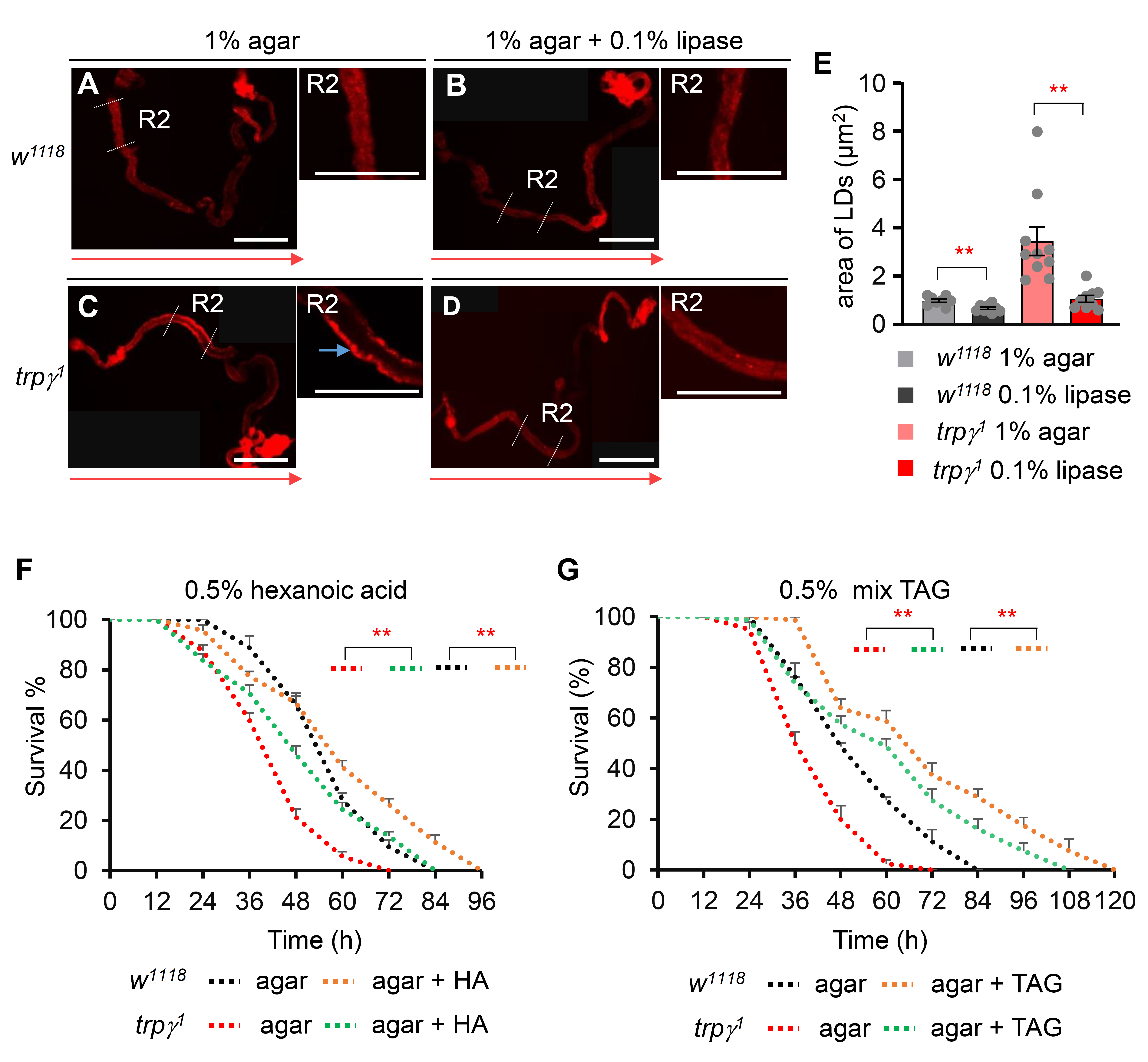

**Figure 6–figure supplement 1:** Measurement of lipase effect in LDs and starvation survival assay after feeding hexanoic acid or mixed TAG**.** (**A**–**D**) Nile red staining of full gut of *w^1118^* and *trpγ^1^* under normal and lipase-fed conditions. (**A** and **B**) Full gut of *w^1118^* under 1% agar fed and 1% agar with 0.1% lipase fed, respectively. (**C** and **D**) Full gut of *trpγ^1^* under 1% agar fed and 1% agar with 0.1% lipase fed, respectively. The scale bar represents 50 µm. Red arrows indicate the orientation of intestine from anterior to posterior. Blue arrow indicates accumulated LDs. (**E**) Measurement of area of LDs in gut of *w^1118^* and *trpγ^1^* after feeding 1% agar or 1% agar with 0.1% lipase (n=3). (**F**) Survival assays to measure the survival time (h) of control (*w^1118^*) and *trpγ^1^* flies after feeding 0.5% hexanoic acid in 1% agarose food (n=4). (**G**) Survival assays of *w^1118^* and *trpγ^1^* flies after feeding 0.5% triglyceride mix (mixture of mono-, di-, tri-) in 1% agar food (n=4).

Means ±SEMs. Single factor ANOVA with Student t-test was used as a *post hoc* test to compare data for **E**. Survival curves were estimated for each group, using a Kaplan-Meier method and compared statistically using the log-rank tests for **F** and **G**. The asterisks indicate significance from the controls (^**^*P* < 0.01).

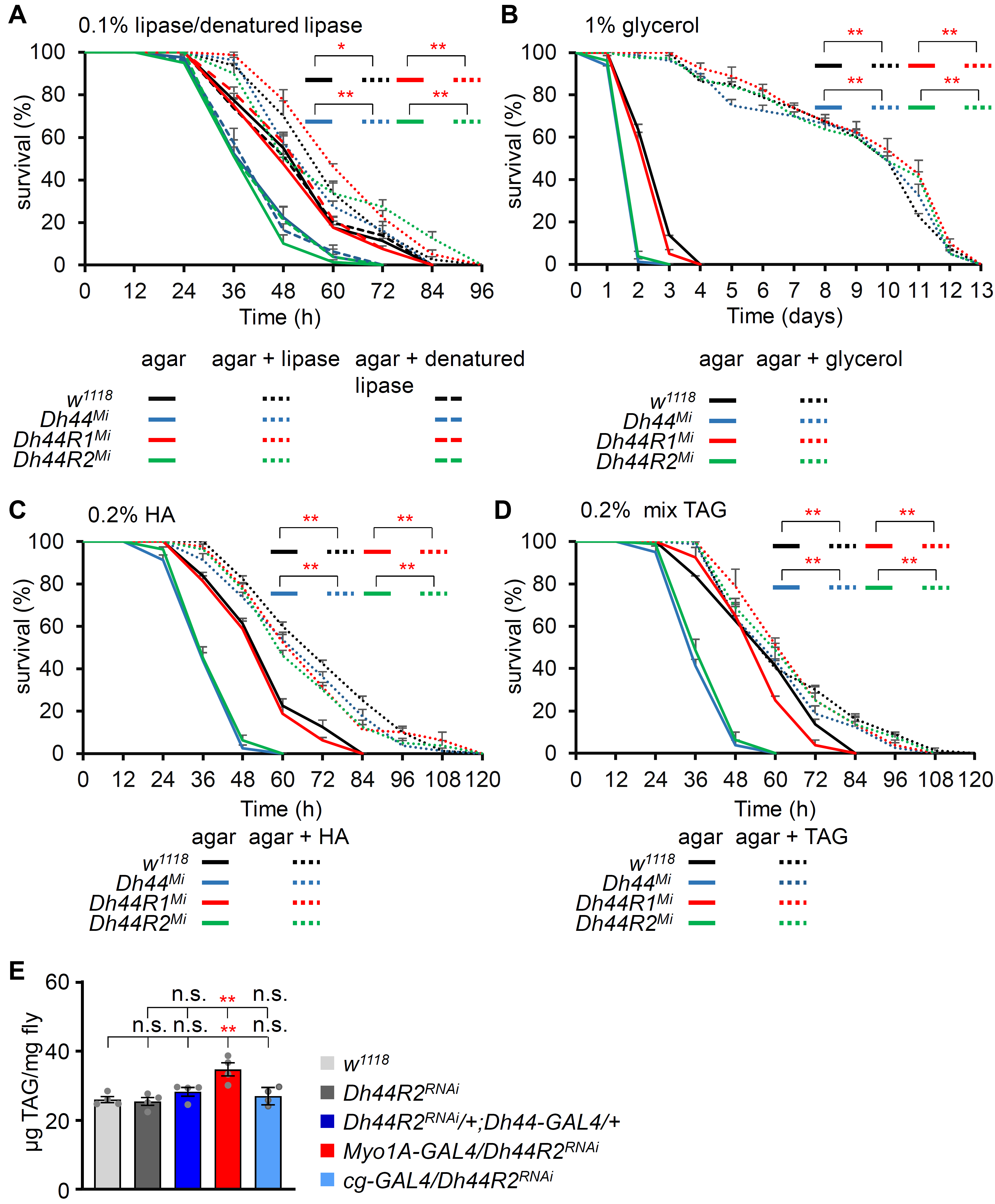

**Figure 7–figure supplement 1:** Measurement of starvation survival time with feeding lipase, glycerol, hexanoic acid and triglyceride mix in 1% agar food. (**A**) Survival assay of *w^1118^*, *Dh44^Mi^*, *Dh44R1^Mi^*, and *Dh44R2^Mi^* after feeding 0.1% lipase and 0.1% denatured lipase in 1% agar food (n=4). (**B**) Survival assay of *w^1118^*, *Dh44^Mi^*, *Dh44R1^Mi^*, and *Dh44R2^Mi^* after feeding 1 % glycerol in 1% agar food (n=4). (**C**) Survival assay of *w^1118^*, *Dh44^Mi^*, *Dh44R1^Mi^*, and *Dh44R2^Mi^* after feeding 0.2 % hexanoic acid in 1% agar food (n=4). (**D**) Survival assay of *w^1118^*, *Dh44^Mi^*, *Dh44R1^Mi^*, and *Dh44R2^Mi^* after feeding 0.2 % triglyceride mix (mixture of mono-, di-, tri-) in 1% agar food (n=4). (**E**) Measurement of TAG level after knockdown of *Dh44R2^RNAi^* in *Dh44*-*GAL4, Myo1A-GAL4,* and *cg-GAL4* (n=4).

Means ±SEMs. Survival curves were estimated for each group, using a Kaplan-Meier method and compared statistically using the log-rank tests. Single factor ANOVA with Scheffe’s analysis was used as a *post hoc* test to compare multiple sets of data in **E**. The asterisks indicate significance from the controls (^*^*P* < 0.05, ^**^*P* < 0.01).
